# Engineering a wolf spider A-family toxin towards increased antimicrobial activity but low toxicity

**DOI:** 10.1101/2024.03.04.583312

**Authors:** Ludwig Dersch, Antonia Stahlhut, Johanna Eichberg, Anne Paas, Kornelia Hardes, Andreas Vilcinskas, Tim Lüddecke

## Abstract

Peptides with insecticidal, antimicrobial and/or cytolytic activities, also known as spider venom antimicrobial peptides (AMPs), can be found in the venoms of RTA-clade spiders. They show translational potential as therapeutic leads. A set of 52 AMPs has been described in the Chinese wolf spider (*Lycosa shansia*), and many have been shown to exhibit antibacterial effects. Here we explored the potential to enhance their antimicrobial activity using bioengineering. We generated a panel of artificial derivatives of an A-family peptide and screened their activity against selected microbial pathogens, vertebrate cells and insects. In several cases, we increased the antimicrobial activity of the derivatives while retaining the low cytotoxicity of the parental molecule. Furthermore, we injected the peptides into adult *Drosophila suzukii* and found no evidence of insecticidal effects, confirming the low levels of toxicity. Our data therefore suggest that spider venom linear peptides can be modified into more potent antimicrobial agents that could help to battle infectious diseases in the future.

## Introduction

Animal venoms provide a remarkably abundant and diverse source of unique proteins and peptides, many with medically-relevant activities [1–3].Venomous animals have recruited many different toxins, which are used to capture prey, to facilitate feeding, for intraspecific competition and to deter predators [4,5]. A particularly rich source of biomolecules is found in spiders which have the potential to yield ∼10 million unique toxins. However, most research thus far has focused on neurotoxic inhibitor cysteine knot (ICK) peptides that act on ion channels in humans and insects, making them suitable as candidates for drugs and bioinsecticides [2,6,7].

More recent analysis of venoms from the wandering RTA-clade spiders have revealed a variety of cytolytic peptides also known as cationic peptides, short linear peptides or spider venom antimicrobial peptides (AMPs) [4,8–14]. They usually contain 15–35 amino acids, form α-helices with a hydrophobic face, and are positively charged. This gives them the ability to attach to various types of biomembranes and permeate them via different modes of action [15–18]. They show potent membranolytic activity in bacteria [19]. The main protective mechanism used by resistant bacteria against conventional antibiotics involves membrane alterations, but AMPs have nonspecific modes of action that bacteria can hardly circumvent [15,20–22]. AMPs are therefore medically relevant antimicrobial agents that could help to turn the tide against the increasing threat of multidrug-resistant bacteria [23].

Here we focus on the Chinese wolf spider (*Lycosa shansia*), formerly known as *Lycosa sinensis*. The analysis of its venom by mass spectrometry revealed 52 AMPs split into eight families known as LS-AMP-A to LS-AMP-H [14]. A recent study focusing on the LS-AMP-A family of 20 α-helical AMPs revealed similarities to the peptides found in frog poison. The LS-AMP-A peptides were found to inhibit bacterial growth but showed no cytotoxicity in assays with Madin-Darby canine kidney (MDCKII) cells [24]. Another study, looking at other *L. shansia* AMPs, reported activity against different clinically relevant drug-resistant bacteria, as well as synergistic effects with traditional antibiotics [25]. This highlights the relevance of RTA-clade spider venom as a promising source of leads for the development of new antimicrobial compounds.

The efficacy of AMPs, including their membranolytic activity, is mainly dependent on their charge and hydrophobicity, which can be modified by amino acid substitutions [26]. The replacement of single amino acids can generate new AMP-like peptides in this seemingly unlimited sequence space [27,28], but rational design based on the bioinformatic prediction of antimicrobial potential may provide a more effective strategy [29]. Here we harnessed our current knowledge on the properties of short linear peptides based on information provided by databases and prediction tools to enhance the antimicrobial potential of known AMPs [15– 17,30]. We produced three derivatives of the natural peptide LS-AMP-A1 by increasing the charge, hydrophobicity or both, and tested them in a broad bioactivity screen. The development of AMPs with enhanced antibacterial activity but minimal activity against mammalian cells and insects will provide promising leads for the development of more effective antimicrobial drugs from spider venom [2,28,31].

## Material and methods

### Peptides

We used the antimicrobial peptide database 3 tool antimicrobial peptide designer [30] to evaluate the natural peptide LS-AMP-A1 for amino acid changes that would influence its charge and hydrophobicity. We manually selected residues for replacement. Accordingly, this led to the design of three engineered derivatives of LS-AMP-A1 which we named LS-AMP-A1_A, LS-AMP-A1_B and LS-AMP-A1_C. We aimed to introduce minimal sequence changes that retained the helical characteristics of the peptide while achieving strong increases in charge, hydrophobicity or both. Therefore, the following modifications were made: In LS-AMP-A1, the serine at position one was modified to isoleucine, in LS-AMP-A1_B we altered the aspartic acid at position seven to lysine and in LS-AMP-A1_C both changes were introduced. The designed peptides were synthesized by a commercial supplier of custom-made peptides (GenScript Biotech, Rijswijk, The Netherlands) via solid phase synthesis as described previously [24].

### Pre-screening growth curve

A cryo-conserved vial of the Gram-negative bacterium *Escherichia coli* (DSM 102053) was transferred to tryptone soy broth (TSB) agar plates and incubated for 24 h at 37 °C. Single colonies were picked and transferred into 3 mL of liquid TSB medium and cultivated overnight at 37 °C, shaking at 180 rpm. Subcultures in fresh liquid medium were prepared, by transferring 40 μL overnight culture into 3 mL fresh media and grown for 4 h at 37 °C before testing. We measured the optical density of each culture at 600 nm (OD_600_) using a DR 2800 Spectralphotometer (Hach Lange) and prepared further dilutions at the preferred OD_600._ The peptides were prepared as 10 mmol/L stocks in DMSO and before each assay they were diluted with TSB medium to 400 μmol/L. TSB medium and 3.85% DMSO were used as negative controls, and 10 μg/mL gentamicin (Sigma-Aldrich) as a positive control for antimicrobial activity. The bacteria were transferred to 96-multiwell plates with volumes of 100 μL per well (50 μL of the bacterial suspension and 50 μL of the test peptide) resulting in final 200 μmol/L peptide concentrations. We prepared growth curves over a 36-h period by taking OD_600_ measurements every 20 min using a BioTek Eon microplate reader. Data were analyzed and visualized using GraphPad Prism v10.0.2.

### Antibacterial activity

A broader screen for antibacterial activity was carried out using a panel of seven bacterial strains as described previously [29]: *Listeria monocytogenes* (DSM 20600), *Micrococcus luteus* (DSM 20030), two *Pseudomonas aeruginosa* strains (DSM 1117 and DSM 50071), *Staphylococcus aureus* (DSM 2569), *Staphylococcus epidermidis* (DSM 28319) and *Escherichia coli* (DSM 102053). We seeded multiple 96-multiwell plates and exposed the panel to 200 μmol/L of each peptide in 100 μL medium, followed by OD_600_ measurements in triplicate using a BioTek Eon microplate reader 18 h after peptide exposure. The growth of each bacterium was normalized to the solvent control cultures in DMSO (100% growth) and medium without bacteria (0% growth). The effect of the engineered peptides was compared to LS-AMP-A1 by applying a paired t-test to show significant differences in growth performance. The data were analyzed and visualized in GraphPad Prism v10.0.2.

### Cell viability assay

The assessment of effects on cell viability was carried out as described earlier [32]. Briefly, Madin-Darby canine kidney II (MDCK II) cells were cultured in Dulbecco’s modified Eagle’s medium (DMEM GlutaMAX) supplemented with 1% penicillin/streptomycin and 10% fetal calf serum (all reagents from Thermo Fisher Scientific, Waltham, MA, USA) and incubated at 37 °C in a 5% CO_2_ atmosphere. The natural peptide LS-AMP-A, its derivatives, and ionomycin (Cayman Chemical, Ann Arbor, MI, USA) as a positive control were dissolved in water to create 10 mmol/L stock solutions. The cells were seeded in 96-well plates and grown to 90% confluency before treatment with 200 μmol/L of the compounds. After incubation for 48 h as above, cell viability was quantified using the CellTiterGlo Luminescent Cell Viability Assay (Promega, Walldorf, Germany). Luminescence was measured in black 96-well plates in a Synergy H4 microplate reader (Biotek, Waldbronn, Germany). Relative light units (RLU) were normalized to the untreated set at 100% and ionomycin as 0%. Means and standard deviations were calculated from triplicate measurements using GraphPad Prism v10.0.2.

### Insecticidal activity

*Drosophila suzukii* flies from a laboratory stock, originally sourced from Ontario, Canada, were reared in a climate chamber at 26 °C with 60% humidity and a 12-h photoperiod. They were fed on a mixture of 10.8% soybean and cornmeal mix, 0.8% agar, 8% malt, 2.2% molasses, 1% nipagin and 0.625% propionic acid. We used adult flies, 4–6 days old, for the insecticidal assay. Each peptide was prepared as a 1 mg/mL solution in distilled water and 46 nL was injected into the thorax using glass capillaries pulled on a P-2000 Laser-Based Micropipette Puller (Sutter Instrument, Novato, CA, USA) held on a Nanoject II device (Drummond Scientific, Broomall, PA, USA). Pure distilled water and 70% ethanol were injected as controls. Injections were performed under a Stemi 508 stereomicroscope (Zeiss). Batches of 10 injected flies were incubated in Ø 29 × 95 mm vials with Ø 30 × 30 mm foam stoppers (Nerbe Plus, Winsen, Germany) filled 1/16 with food medium. Survival rates were assessed 1, 3 and 24 h post-injection. Data were visualized using GraphPad Prism v10.0.2

## Results

### Sequence alterations in engineered peptides and their physicochemical consequences

Our predictions suggested that three derivatives of the natural peptide LS-AMP-A1 with a maximum of two amino acid substitutions and no change in length would increase its activity. The 11-amino-acid motif LVKSKVGQTGLL at the C-terminus of all derivatives remained the same. In the first derivative (LS-AMP-A1_A), the serine at position 1 was replaced with the hydrophobic residue isoleucine resulting in a 10% increase in hydrophobicity. In the second derivative (LS-AMP-A1_B), the aspartic acid residue at position 7 was replaced with lysine, thus leading to an overall +2 increase in charge compared to the natural variant. The final derivative (LS-AMP-A_C) featured both of the modifications described above, increasing the total hydrophobicity to 70% and the total charge to +3.

**Table 1.** Physiochemical properties of LS-AMP-A1 and its derivatives. Given are the name of each synthesized peptide, its sequence plus number of amino acids and the calculated hydrophobicity and charge.

| Name | Sequence | #AAs | Hydrophobicity | Charge |
| --- | --- | --- | --- | --- |
| LS-AMP-A1 | SLFGLLDLVKSKVGQTGLL | 19 | 61 % | +1 |
| LS-AMP-A1_A | ILFGLLDLVKSKVGQTGLL | 19 | 71 % | +1 |
| LS-AMP-A1_B | SLFGLLKLVKSKVGQTGLL | 19 | 60 % | +3 |
| LS-AMP-A1_C | ILFGLLKLVKSKVGQTGLL | 19 | 70 % | +3 |

### Pre-screening of antimicrobial activity against *E*. *coli*

If the bioinformatically designed peptides with altered hydrophobicity and charge retain their natural antimicrobial activity is investigated in a pre-screening experiment. We tested the four peptides for general antibacterial properties against *E. coli*. The maximum growth rate was marginally reduced in the presence of the natural peptide, but the effect was much more potent in the presence of the derivatives. All three of the derivative peptides reduced the growth rate of the bacteria by 50% compared to the natural peptide. These promising results prompted us to carry out further screenings against a broader array of bacteria.

### Antibacterial activity against a wider range of strains

In the broader screen, we assessed the inhibitory activities of the peptides against seven different bacterial strains including several clinically relevant ones (Figure 2). The natural peptide LS-AMP-A1 did not significantly reduce the growth of any of these strains. In contrast, LS-AMP-A1_C caused a significant reduction in the growth of *P. aeruginosa* (1117), *M. luteus*, and *E. coli*, but had no significant effect on the other four strains, in comparison to the natural peptide. LS-AMP-A1_B significantly reduced the growth of *S. aureus* and *E. coli* while also inhibiting *P. aeruginosa* (50071), *M. luteus* and *L. monocytogenes*, albeit not to a statistically significant extent. This derivative had no significant effect against *P. aeruginosa* (1117) or *S. aureus*. Finally, LS-AMP-A1_A had no significant effects against any of the test strains. Overall, no peptide had visible effects against either *S. epidermidis* or *P. aeruginosa* (50071).

**Figure 1.**
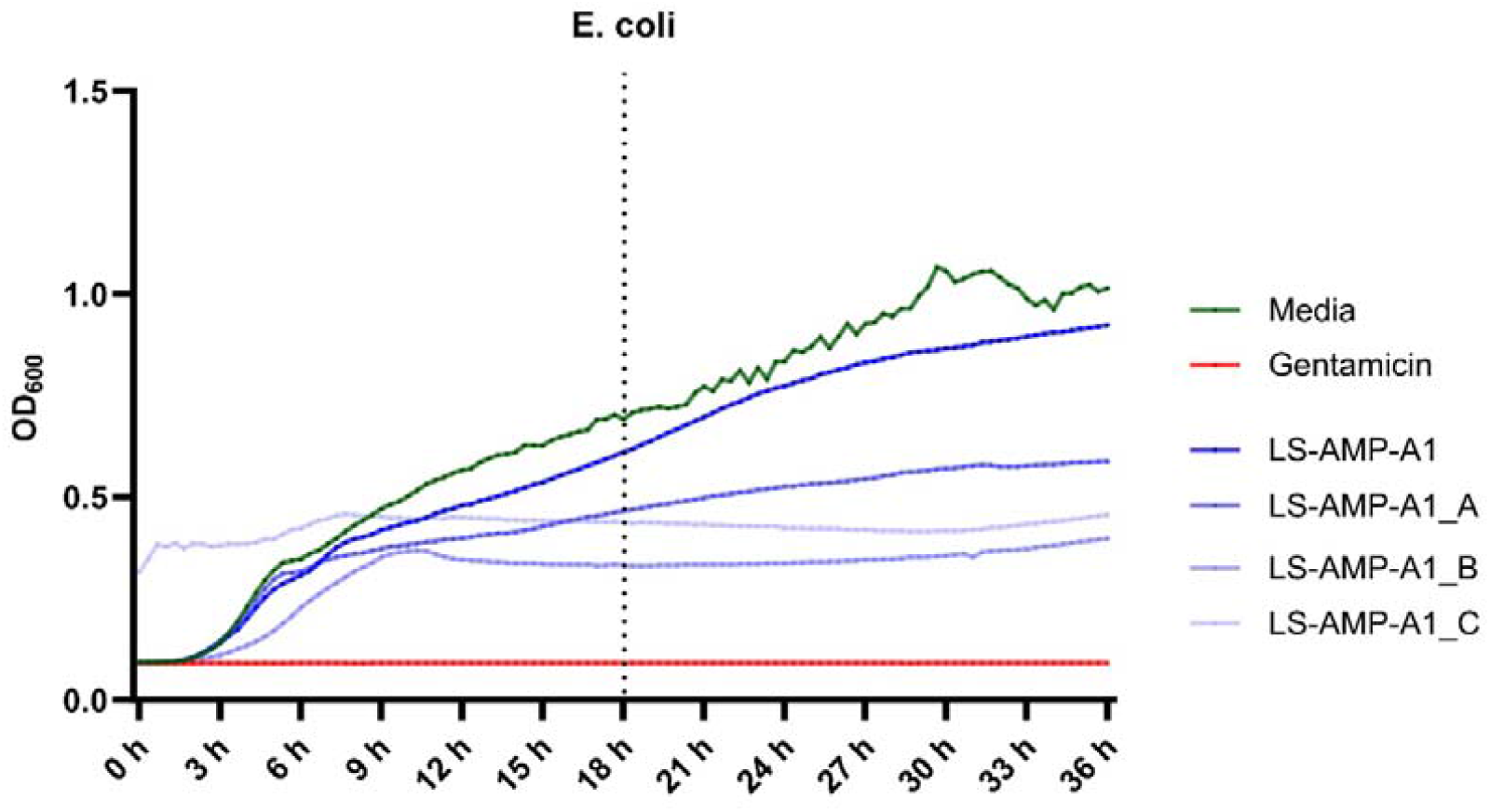
Growth curve of *E. coli* in presence of LS-AMP-A1, its derivatives, and Gentamicin over 36 hours with measurements every 20 min. The dotted vertical line indicates the 18-hour time point.

**Figure 2.**
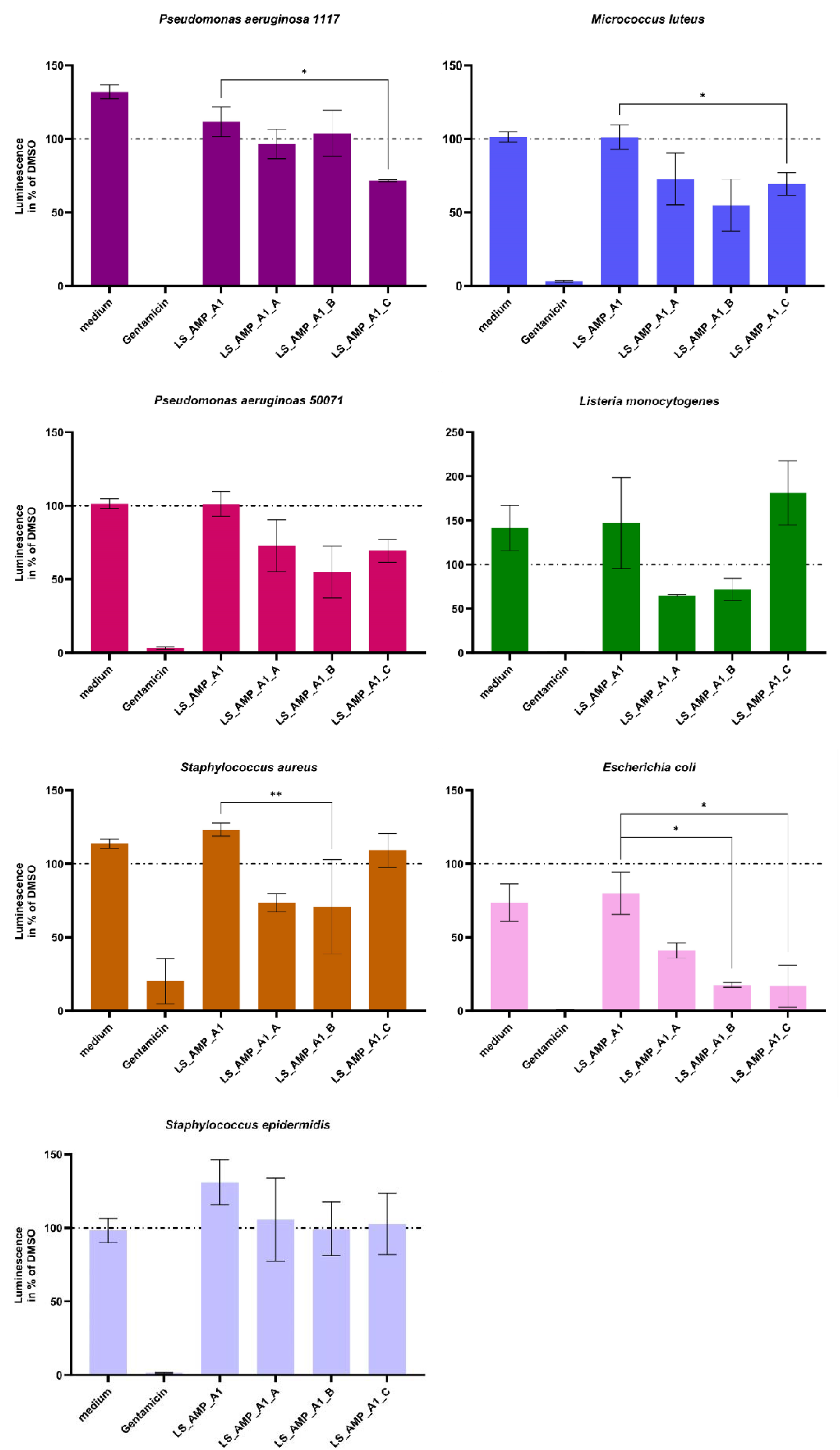
Activity of LS-AMP-A1 and its derivatives on the growth of seven different bacterial strains after 18 hours. Significant differences between the natural peptide and its derivatives evaluated by paired t-test and indicated with asterisk (p ≤ 0,05 *, p ≤ 0,01 **, p ≤ 0,001 ***). Data are means with error bars (n=3).

### Cell viability Assay

Given the membranolytic potential of AMPs ([19,33]), we tested the four peptides for cytotoxicity against the mammalian cell line MDCK II at a concentration of 200 μM. Only the peptide LS-AMP-A_C showed cytotoxic effects at this concentration (Figure 3).

**Figure 3.**
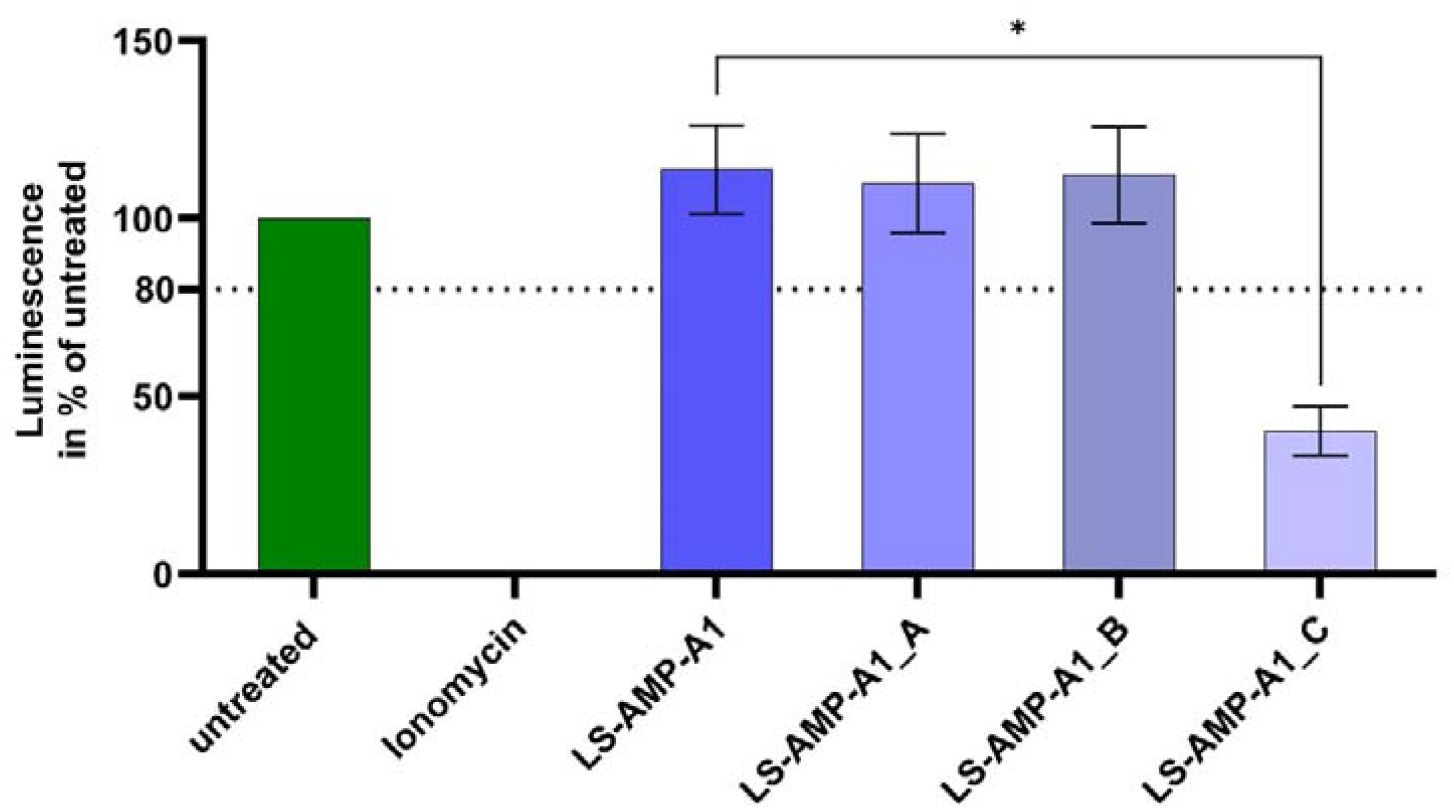
Cell viability of engineered and natural peptides. Cytotoxic activity of LS-AMP-A1 and its derivatives on MDCKII cells after 48h of treatment with each peptide. Luminescence values normalized to untreated as 100% and Ionomycin as 0%. Data are means with error bars (n=3) evaluated by paired t-test and indicated with asterisk (p ≤ 0,05 *, p ≤ 0,01 **, p ≤ 0,001 ***).

**Figure 4.**
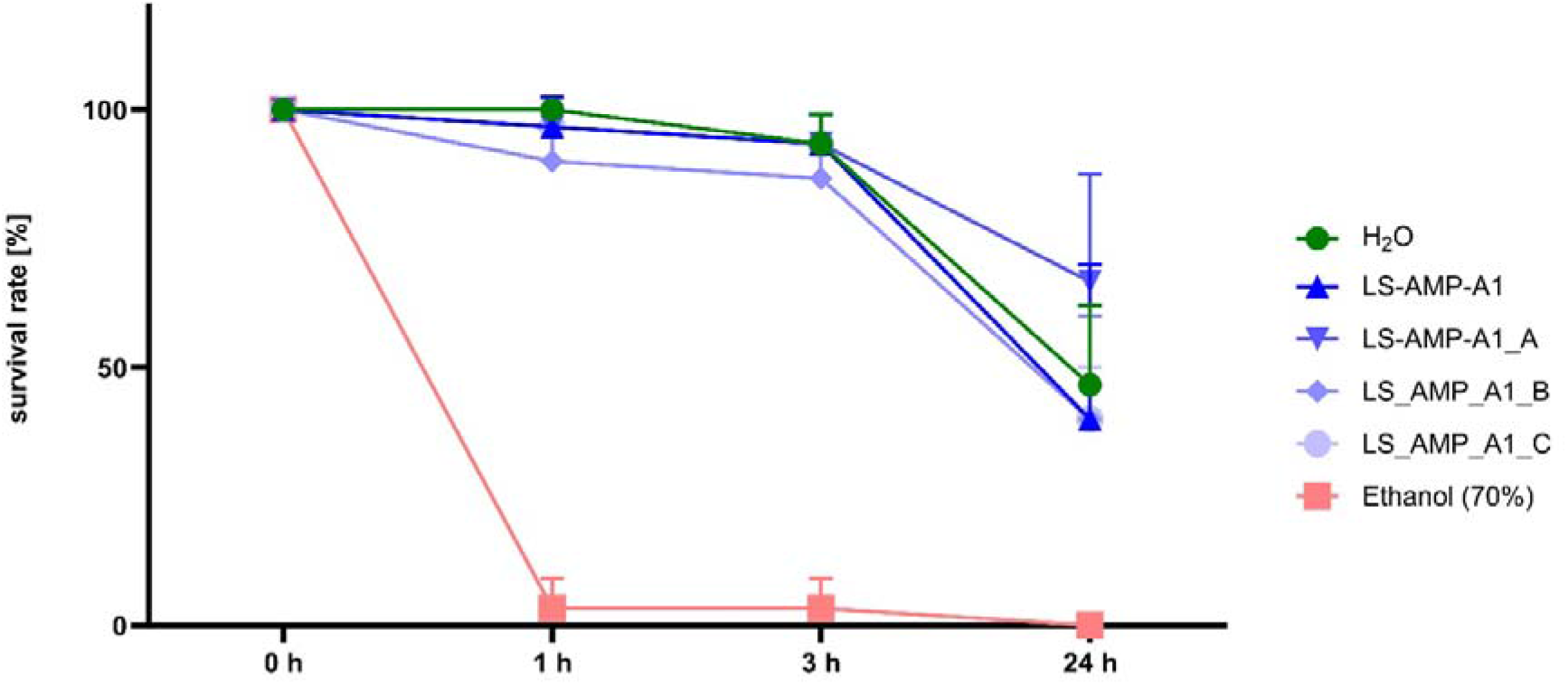
Insecticidal activity of LS-AMP-A1 and its derivatives against *Drosophila suzukii* flies. Immediate effects 1h and 3h after injection were assessed, as well as effects 24h hours post injection. 1 mg/mL peptide solution, H2O and Ethanol were injected in volumes of 46 nL in 3 sets of 10 animals each.

### Insecticidal Activity

Finally, we injected the peptides into the thorax of *Drosophila suzukii* adult flies to determine their insecticidal activity as an approximation for off-target effects in the environment. Given that spider toxins are expected to show immediate effects on insects, we screened the effects for 24 h post-injection, after 1, 3 and 24 h. None of the peptides showed direct lethal effects 1 or 3 h after injections, and there was no evidence of enhanced lethality compared to water-injected controls after 24 h.

## Discussion

The hydrophobicity and charge distribution of natural AMPs are the main properties that determine their biological activity, therefore informing the design of novel synthetic AMPs derived from natural templates [19,26,27,33]. Accordingly, we focused on these two physicochemical properties to enhance the activity of known AMPs from the Chinese wolf spider *L. shansia*. We selected LS-AMP-A1 as our first template peptide because it showed promising but low levels of antimicrobial activity in preliminary screens yet exhibited no cytotoxicity [24]. The application of bioinformatic tools facilitated the rational design of enhanced derivatives by identifying specific residues that could be replaced to increase the charge and/or hydrophobicity of the natural peptide.

Short linear peptides from spider venoms are thought to promote the uptake of co-injected venom components, suggesting trophic functions or roles to deter predators, and this has been confirmed for several peptides [4,34]. However, the recent analysis of four LS-AMP families does not support such trophic functions, and suggests instead the principal role may be to suppress the growth of bacteria in the venom glands [24,25]. Our results support this hypothesis because we found no evidence of cytotoxicity or insecticidal activity when testing the natural peptide LS-AMP-A1 and the derivatives LS-AMP-A1_A and LS-AMP-A1_B, while LS-AMP-A1_C had increased cytotoxicity. However, all three derivatives retained their antimicrobial activity and even showed a significant increase in potency against several bacteria.

LS-AMP-A1_A did not cause a statistically significant reduction in the growth of the bacterial strains we tested, but we observed slight reductions in the growth of *M. luteus* and *S. aureus*. This suggests that a single amino-acid exchange that increases hydrophobicity is not sufficient to cause a marked increase in antibacterial activity. In contrast, LS-AMP-A1_B retained the hydrophobicity of the natural peptide but gained a higher charge due to the introduction of a lysine residue. This increased its antibacterial activity significantly against Gram-positive *S. aureus* and Gram-negative *E*. coli, additionally inhibiting *P. aeruginosa* (50071), *M. luteus* and *L. monocytogenes*, albeit not to a statistically significant extent. This result indicates that substitutions that increase the charge of LS-AMP-A1 are more likely to influence the antibacterial activity than changes in hydrophobicity. Finally, LS-AMP-A1_C combined the modifications of charge and hydrophobicity that were separately introduced into the other two derivatives. This variant significantly reduced the growth of three bacterial strains, namely Gram-negative *P. aeruginosa* (1117), Gram-positive *M. luteus*, and Gram-negative *E. coli*, thus representing the most effective modifications among the three peptides, but also shows increased cytotoxicity. Our data therefore shows that the activity of peptides can be increased by single amino acid exchanges. Increasing the hydrophobicity alone is not sufficient to modify the activity of the native LS-AMP, but simultaneously modifying the hydrophobicity and charge has a more potent effect than modifying the charge alone.

The ability to inhibit both Gram-positive and Gram-negative bacteria shows that the activity of the peptides is not limited by the membrane characteristics of the bacteria. However, the absence of cytotoxicity and insecticidal activity for derivates A and B implies that any interactions with mammalian or insect cells do not affect cell viability. The cytotoxic and insecticidal effects of some peptides depend on interaction partners, with which they synergise [15,35]. It may therefore be possible for our modified peptides to inhibit bacterial growth even more by combining them with other venom components, or even with conventional antibiotics [25]. The absence of insecticidal effects, supporting the role of LS-AMPs as an antimicrobial defense peptide to prevent the colonization of venom glands by pathogens, will allow the development of derivative peptides as ecologically benign antimicrobials.

In conclusion, three different derivates of LS-AMP-A1 with differing activity profiles have been generated. Our study shows that the antimicrobial activity of known spider venom AMPs can be enhanced in different directions by simple amino acid substitutions, highlighting their value as templates for bioengineering. Nature provides a rich and largely untapped reservoir of spider toxins with a range of bioactivities, and the ability to enhance or otherwise modify them by replacing individual amino acids or groups provides an exponentially larger range of molecules for testing and development into therapeutics to combat the threat of multi-resistant bacteria [2]. Rather than abandoning natural AMPs with weak antibacterial effects in favor for more potent alternatives, it may therefore be more productive to consider modifications that increase charge and hydrophobicity to gain increased activities, as demonstrated here for LS-AMP-A1_C. Although considering only a single parental AMP, we have shown that the replacement of a single amino acid has the potential to significantly improve antimicrobial potency without inducing cytotoxicity against mammalian cells or insects. Overall, our results provide additional support for the importance of animal venom toxins as sources of biomedical innovation and underpin the importance of bioengineering natural components towards improved activity [36].

## Supporting information

Raw Data Insecticidal Assay

Raw Data Bacteria OD

Raw Data Antimicrobial Assay

Raw Data Cytotoxicity Assay

Raw Data Growthcurve

## Notes

### Competing Interest Statement

The authors have declared no competing interest.

