## Supplementary material for "Engineering a wolf spider A-family toxin towards increased antimicrobial activity but low toxicity": Raw Data Bacteria OD

| **Bacteria strains** | **Preferred**  **Assay OD_600_** |
| --- | --- |
| *L. monocytogenes* | 0.00125 |
| *M. luteus* | 0.005 |
| *P.aeruginosa* (1117) | 0.005 |
| *P.aeruginosa* (50071) | 0.00125 |
| *S. aureus* | 0.00125 |
| *S. epidermidis* | 0.000625 |
| *E. coli* | 0.000312 |
